# Unliganded and CMP-Neu5Ac bound structures of human α-2,6-sialyltransferase ST6Gal I at high resolution

**DOI:** 10.1101/2020.06.07.138503

**Authors:** Deborah Harrus, Anne Harduin-Lepers, Tuomo Glumoff

## Abstract

Sialic acid residues found as terminal monosaccharides in various types of glycan chains in cell surface glycoproteins and glycolipids have been identified as important contributors of cell-cell interactions in normal vs. abnormal cellular behavior and are pivotal in diseases such as cancers. In vertebrates, sialic acids are attached to glycan chains by a conserved subset of sialyltransferases with different enzymatic and substrate specificities. ST6Gal I is a sialyltransferase using activated CMP-sialic acids as donor substrates to catalyze the formation of a α2,6-glycosidic bond between the sialic acid residue and the acceptor disaccharide LacNAc. Understanding sialyltransferases at the molecular and structural level shed light into the function. We present here two human ST6Gal I structures, which show for the first time the enzyme in the unliganded state and with the full donor substrate CMP-Neu5Ac bound. Comparison of these structures reveal flexibility of the catalytic loop, since in the unliganded structure Tyr354 adopts a conformation seen also as an alternate conformation in the substrate bound structure. CMP-Neu5Ac is bound with the side chain at C-5 of the sugar residue directed towards empty space at the surface of the protein. Furthermore, the exact binding mode of the sialic acid moiety of the substrate directly involves sialylmotifs L, S and III and positions the sialylmotif VS in the immediate vicinity.

**PROTEIN DATA BANK ACCESSION CODES:** Atomic coordinates and structure factors of the human wild-type unliganded and CMP-Neu5Ac bound ST6Gal I have been deposited with the PDB with accession codes 6QVS and 6QVT, respectively.

## 1. INTRODUCTION

Sialylation is defined as occurrence of sialic acids as terminal sugar residues in glycolipids and in hybrid- and complex type N- or O-glycans of cell surface glycoproteins (Zhang *et al*., 2018; Varki *et al*., 2017). Sialic acids are monosaccharides with a 9-carbon backbone containing a carboxylic acid group, an exocyclic side chain, and an N-acetyl group. The most common sialic acid residue in human tissues is the *N*-acetylneuraminic acid (5-acetamido-3,5-dideoxy-D-glycero-D-galacto-non-2-ulosonic acid, Neu5Ac). Other naturally occurring sialic acids contain various *N*- or *O*- linked groups (e.g. *N*-glycolyl, *O*-methyl or *O*-acetyl) as substituents (Schauer & Kamerling, 2018). These unusual features in a monosaccharide residue combined with their exposed position in the glycan chains let it presume that they give cells specific functional properties. Indeed, it is commonly known that sialylation is an important determinant in the role of glycans in e.g. cell-cell recognition and interaction. Therefore not surprisingly, abnormal sialylation is associated with several groups of diseases among which are cancers, where correct interactions between cells are pivotal (Varki, 2017).

Sialic acid residues are added to glycan chains by sialyltransferases (ST) from an activated nucleotide-sugar donor substrates, the Cytidine-5′-monophosphate sialic acid (CMP-sialic acid). Like all glycosyltransferases (GTs), STs are also grouped into GT-families based on enzyme activity, specificity and modular nature in the Carbohydrate-Active enZYmes (CAZy) database (www.cazy.org; Lombard *et al*., 2014). There are currently 110 CAZy GT-families of which 5 comprise of different STs: family GT29 contains eukaryotic, viral and some bacterial STs, while families GT38, GT42, GT52 and GT80 comprise of bacterial STs (Audry *et al*., 2011; Petit *et al*., 2018). In the GT29, vertebrate STs are further classified based on their distinct enzyme activity, glycosidic linkage formed (α2,3-, α2,6- and α2,8-linkages) and monosaccharide acceptor: ST3Gal, ST6Gal, ST6GalNAc and ST8Sia (Harduin-Lepers, 2010; Harduin-Lepers *et al*., 2005). ST6Gal I (β-galactoside α-2,6-sialyltransferase I (EC 2.4.99.1)), the subject of this study, belongs to GT29. It connects a galactose C6 OH-group and a sialic acid C2 OH-group into an O-glycosidic bond in α-configuration. In human tissues, ST6Gal I uses primarily the Cytidine-5′-monophospho-*N*- acetylneuraminic acid (CMP-Neu5Ac) as a donor substrate in the reaction in humans is CMP-β- Neu5Ac.

Sialyltransferases are found in the trans-Golgi network membranes as type II membrane proteins with a short N-terminal portion on the cytosolic side, a single transmembrane α-helix, a presumably disordered stem region and a catalytic domain on the luminal side (Audry *et al*., 2011). Sequence analyses suggested, and subsequent structural work confirmed, the presence of four conserved peptide motifs in the catalytic domain of STs named sialylmotif L (for Large), S (Small), III (3^rd^) and VS (Very Small) (Jeanneau *et al*., 2004; Datta, 2009; Audry *et al*., 2011; Patel & Balaji, 2006; Harduin-Lepers, 2010). Site directed mutagenesis studies confirmed the implication of these sialylmotifs in the donor and acceptor binding sites and catalysis (Datta and Paulson 1995; Datta 2009). Very few crystal structures of mammalian sialyltransferases are known (Table 1) and they all represent the soluble catalytic domain with varying small stretch of the stem domain included. The first structure - pig ST3Gal I - was published by Rao and coworkers in 2009 (Rao *et al*., 2009), followed by structures of rat (Meng *et al*., 2013) and human (Kuhn *et al*., 2013) ST6Gal I as well as human ST8Sia III (Volkers *et al*., 2015). The most recent structure is that of human ST6GalNAc II (Moremen *et al*., 2018). All these ST structures belong to the same fold type, named GT-A (variant 2). Important for assessment of structural details, especially when comparing structures with different ligands bound, is that sialylmotif L is engaged mainly in binding the donor substrate, while sialylmotifs S, III and VS are engaged in binding the acceptor substrate or both substrates. In addition, multiple sequence alignment of vertebrate sialyltransferases revealed the existence of family motifs specific of each sialyltransferase family i.e. ST3Gal, ST6Gal, ST6GalNAc and ST8Sia. These family motifs named “a”, “b”, “c”, “d” and “e” correspond to another level of amino acid conservation that could be relevant for linkage specificity and monosaccharide acceptor specificity (Harduin-Lepers 2010, Patel and Balaji 2006)

**Table 1.** Sialyltransferase crystal structures in the Protein Data Bank. Each structural entry is characterized by organism, enzyme acronym, amino acid range of the polypeptide seen in the crystal structure, possible bound ligand, resolution (in Å), PDB access code and literature reference. Abbreviations of the ligands are: **A2G**, N-acetyl-2-deoxy-2-amino-galactose; **BMA**, β-D-mannose; **C or C5P**, cytidine-5’-monophosphate; **CDP**, cytidine-5’-diphosphate; **CMP-Neu5Ac**, cytidine-5′-monophospho-N-acetylneuraminic acid**; CSF**, cytidine-5’-diphosphate-3-fluoro-N-acetyl-neuraminic acid; **CTN**, 4-amino-1-β-D-ribofyranosyl-2(1H)-pyrimidinone; **CTP**, cytidine-5’-triphosphate; **FUC**, α-fucose; **FUL**, β-L-fucose; **GAL**, β-D-galactose; **MAN**, α-D-mannose; **NAG**, N-acetyl-D-glucosamine; **NGS**, 2-(acetylamino)-2-deoxy-6-O-sulfo-β-D-glucopyranose; **SIA**, O-sialic acid.

| Organism | Enzyme | AA range | Ligand | Resolution | PDB id | Ref |
| --- | --- | --- | --- | --- | --- | --- |
| Pig | ST3Gal I | 46-343 | - | 1.90 | 2WML | Rao 2009 |
| Pig | ST3Gal I | 46-343 | A2G, C5P, GAL | 1.55 | 2WNB | Rao 2009 |
| Pig | ST3Gal I | 46-343 | A2G, GAL | 1.25 | 2WNF | Rao 2009 |
| Human | ST6Gal I | 89-406 | BMA, CTN, GAL, MAN, NAG | 2.09 | 4JS1 | Kuhn 2013 |
| Human | ST6Gal I | 89-406 | BMA, C, FUL, GAL, MAN, NAG | 2.30 | 4JS2 | Kuhn 2013 |
| Human | ST6Gal I | 132-406 | - | 1.60 | 6QVS | present study |
| Human | ST6Gal I | 132-406 | CMP-Neu5Ac | 1.70 | 6QVT | present study |
| Rat | ST6Gal I | 95-403 | NAG | 2.40 | 4MPS | Meng<br>2013 |
| Human | ST6GalNAc2 | 66-374 | NAG | 3.10 | 6APJ | Moremen<br>2018 |
| Human | ST6GalNAc2 | 66-374 | C5P, NAG | 2.35 | 6APL | Moremen<br>2018 |
| Human | ST8Sia III | 81-380 | BMA, CDP,<br>FUC, NAG | 2.07 | 5BO6 | Volkers<br>2015 |
| Human | ST8Sia III | 81-380 | FUC, CTP,<br>NAG | 1.85 | 5BO7 | Volkers<br>2015 |
| Human | ST8Sia III | 81-380 | MAN, FUC,<br>BMA, NAG | 2.70 | 5BO8 | Volkers<br>2015 |
| Human | ST8Sia III | 81-380 | NGS, FUC,<br>GAL, CSF,<br>NAG, SIA | 2.30 | 5BO9 | Volkers<br>2015 |
| Human | ST8Sia III | 81-380 | NAG, BMA,<br>FUC | 2.15 | 5CXY | Volkers to<br>be<br>published |

In the present work, we have solved the crystal structure of human ST6Gal I. We also compare this structure with human and rat ST6Gal I (Kuhn *et al*. 2013; Meng *et al*., 2013), pig ST3Gal I (Rao *et al*., 2009) and human ST6GalNAc II (Moremen *et al*. 2018). Our results add to the structural knowledge of sialyltransferases in at least three important ways: (i) we present the first unliganded structure of an ST6Gal sialyltransferase; (ii) we present the first ST structure with the natural donor substrate CMP-Neu5Ac bound and seen in its entirety as part of the liganded structure; and (iii) these structures are at clearly highest resolution so far for an ST6Gal sialyltransferase.

## 2. MATERIALS & METHODS

### 2.1. Protein expression and purification

The catalytic domain of the human ST6Gal I (residue range 132-406) was expressed in a BL21(DE3) *Escherichia coli* strain (Invitrogen), using a polycistronic Ptac vector containing the CyDisCo system (Gąciarz *et al*. 2017) enabling disulfide bond formation in the cytoplasm of *E. coli*. Protein expression was induced overnight at 30°C with 1 mM Isopropyl-β-D-thiogalactopyranoside (IPTG). Bacteria were pelleted and lysed by sonication in lysis buffer (50 mM Sodium Phosphate, pH 7.2). Proteins were purified with a Bio-Scale Mini Profinity Nickel cartridge (Bio-Rad) followed by a Superdex 200 (GE Healthcare) and concentrated in the final buffer (20 mM Sodium Phosphate, pH 7.2, 150 mM NaCl). The purity was assessed >95% by SDS-PAGE.

### 2.2. Crystallization, ligand soaking and data collection

Crystals were obtained by the hanging drop vapor diffusion method at room temperature. Equal volumes of protein solution (5.6 mg/mL in 20 mM sodium phosphate, pH 7.2, 150 mM NaCl) and well solution (0.1 M Bis-Tris pH 5.5, 0.1 M Ammonium Acetate, 8% PEG 20.000 or 10% PEG 6000) were mixed. Since the crystals were easily breakable, the following procedure has been used for soaking with CMP-Neu5Ac (CMP-N-acetylneuraminic acid sodium salt (MC04391, Carbosynth Limited, UK), in order to reduce the need for crystal manipulation: skin was removed from the top of the drop, then 0.2 uL of 100 mM CMP-Neu5Ac solution was added directly on top of a 2 uL drop, leading to a final concentration of CMP-Neu5Ac of 10 mM in the drop. Different soaking times were tried, ranging from 1 hour to 6 days. Crystals were harvested and soaked for a few seconds in a cryo-solution (crystallization solution supplemented with 20% glycerol, as well as 10 mM CMP-Neu5Ac in case of the CMP-Neu5Ac bound crystals) and flash-cooled in liquid nitrogen. Several data sets were collected and processed at the beamlines ID30A-3 and ID23-1 of the European Synchrotron Radiation Facility (ESRF), Grenoble, France, as remote sessions, and two high-resolution data sets including one with the best ligand occupancy selected for refinement (Table 2).

**Table 2.** Data collection and refinement statistics. Statistics for the highest-resolution shell are shown in parentheses.

|  | ST6Gal I unliganded (6QVS) | ST6Gal I with CMP-Neu5Ac<br>(6QVT) |
| --- | --- | --- |
| Space group | C 1 2 1 | C 1 2 1 |
| Cell parameters (Å, °) | 138.816 94.369 64.472 90 94.309<br>90 | 136.661 93.921 63.718 90 94.389<br>90 |
| Resolution range (Å) | 38.04 - 1.60 (1.66 - 1.60) | 46.96 - 1.70 (1.76 - 1.70) |
| Molecules per asymmetric unit | 2 | 2 |
| $V_m$ (Å<sup>3</sup>/D) | 3.20 | 3.10 |
| Mean I/sigma(I) | 17.43 (2.28) | 16.36 (1.92) |
| Completeness (%) | 98.86 (99.09) | 98.93 (98.40) |
| Unique reflections | 107819 (10784) | 87229 (8679) |
| Multiplicity | 5.0 (5.1) | 3.5 (3.5) |
| RMS bonds(Å)/angles(°) | 0.009/1.04 | 0.006/0.82 |
| R-work/ R-free | 0.1811/0.2005 (0.2447/0.2568) | 0.1817/0.2042 (0.2773/0.3086) |
| Wilson B-factor | 22.91 | 24.09 |
| Number of atoms (non-H) | 4886 | 4835 |
| protein | 4457 | 4563 |
| solvent | 409 | 258 |
| ligands | 20 | 14 |
| Ramachandran favoured (%) | 98.1 | 97.7 |
| Ramachandran allowed (%) | 1.9 | 2.3 |

### 2.3. Structure determination and refinement

Data were processed with the XDS package (Kabsch, 2010a; Kabsch, 2010b). Previously published crystal structure of the human ST6Gal I was used as the search model for molecular replacement (Protein Data Bank (PDB) code 4JS2) (Meng *et al*., 2013). Molecular replacement was performed with Phaser (McCoy *et al*., 2007). Models were rebuilt with COOT (Emsley *et al*., 2010) and refined with phenix.refine (Adams *et al*., 2010). Structural figures were generated using the PyMOL Molecular Graphics System, Version 1.3 Schrödinger, LLC.

## 3. RESULTS AND DISCUSSION

### 3.1. General description and quality of the structures

In the present work we used the wild-type human ST6Gal I as a construct containing residues 132-406 due to inability to express in sufficient amount, to purify and to crystallize longer constructs. The construct is lacking a 44-residue “neck” region, which includes the end of the stem domain and the beginning of the catalytic domain. Prism-shaped plates with maximum dimensions of ca. 100 micrometers grew within three to seven days, and diffracted to 1.6 Å resolution. Data collection and refinement statistics are presented in Table 2. Both the unliganded (PDB 6QVS) and the CMP-Neu5Ac liganded (PDB 6QVT) protein crystallized with two molecules per asymmetric unit. The two molecules (chain A and chain B) appear not to represent a biologically significant dimer based on jsPISA (Krissinel, 2015) parameters, including a 651 Å^2^ interface area, and gel filtration (not shown). In dynamic light scattering (DLS) we observed fast equilibrium between monomers and higher oligomers.

### 3.2. The unliganded structure of ST6Gal I

For the two molecules in the asymmetric unit the polypeptide chain could be built for residues 132-406 (both chains), except for residues 367-370 in chain A, where tracing was not possible due to no clear electron density. Residues 366-370 in chain B have been built using a low contour level, yet the electron density is weak and fragmented at 1 sigma level. The two chains are very similar with an RMS value of 0.204 Å. In both molecules there are disulfide bridges between cysteines 184-335, 353-364 and 142-406.

### 3.3. The CMP-Neu5Ac bound structure of ST6Gal I

The best ligand occupancy after data collection was observed in the crystals soaked with CMP-Neu5Ac for 3h. The crystals soaked for 24h or more showed the same ligand occupancy but resulted in the phosphate moiety to be hydrolyzed (data not shown). There are tiny features of positive and negative electron density visible at the phosphate of the CMP-Neu5Ac in the presented structure, indicating the essentiality of a short soaking time to capture the ligand. It is known that CMP-Neu5Ac undergoes a relatively rapid hydrolysis under acidic conditions, such as the conditions that gave the crystals (pH 5.5): the carboxylic acid attacks the phosphate group, and easily hydrolyses to CMP and Neu5Ac (Kajihara *et al*., 2011).

The liganded structure contains the donor substrate in full. Otherwise the building and refinement produced a very similar structure with the unliganded enzyme: in chain A residues 365-371 and in chain B residues 366-371 could not be traced. This region corresponds in part to ST family 29 motif “d” (Patel & Balaji, 2006; Harduin-Lepers 2010; Audry *et al*., 2011), which is a ST6Gal family-motif supplementing the general L, S, III & VS sialylmotifs existing in all sialyltransferases, and could be engaged in acceptor substrate binding. In chain B the first two residues of the construct at positions 132 and 133 are also disordered. The two chains are very similar with an RMS value of 0.211 Å. The occupancy of the CMP-Neu5Ac was refined (fixed for all atoms in residues) to 87% in chain A and 91% in chain B. A detailed description of the binding mode of the Neu5Ac moiety of the CMP-Neu5Ac substrate - as seen for the first time - is given in Table 3 and in Figure 1.

**Table 3.** Interacting atoms and types of interaction between ST6Gal I and the sialic acid moiety. It is noteworthy that sialyl motifs L, S and III are directly involved in binding, and motif VS being in the immediate vicinity of the sialic acid moiety. See also Figure 1d.

| ST6 residue and atom | Sialic acid atoms | Type of interaction | motif * |
| --- | --- | --- | --- |
| Asn212 ND2 | O1A, O6 | Hydrogen bond | Sialylmotif L |
| Asn233 ND2 | O1A, O6, O7 | Hydrogen bond | Family motif a,<br>immediately after L |
| Gln235 NE2 | O7, O10 | Hydrogen bond | Family motif a,<br>immediately after L |
| Ser323 HG, N | O1A, O1B | Hydrogen bond | Sialylmotif S |
| Tyr354 OH | N5 | Hydrogen bond | Sialylmotif III |
| Ala363 CB | C18 | Short hydrophobic int. | immediately before<br>VS, family motif d |

\* Based on conserved motifs from sequence comparison of GT-A structural superfamily (Audry *et al.*, 2011)

**Figure 1.**
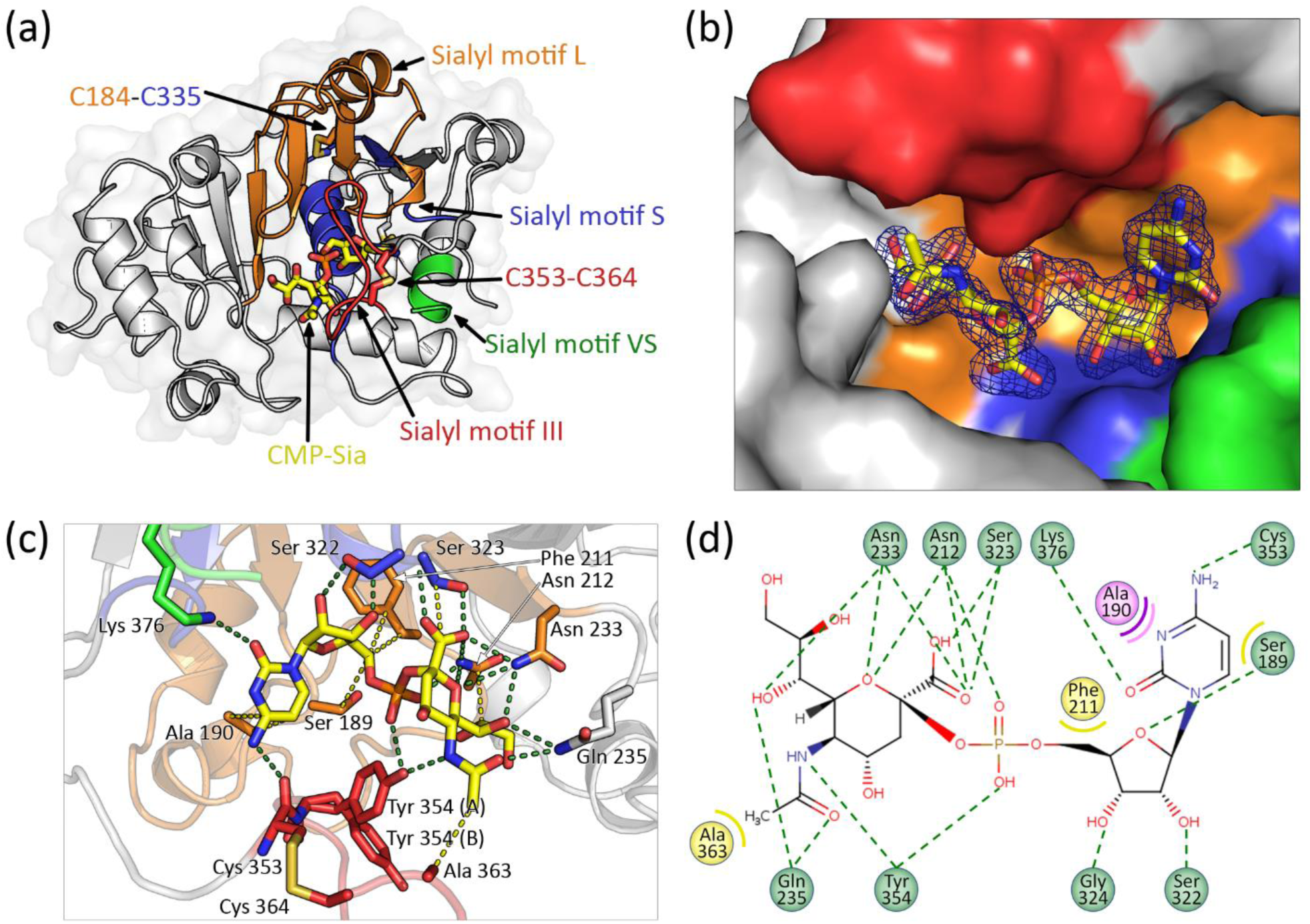
Structure of the human ST6Gal I in the CMP-Neu5Ac bound form. (a) Cartoon view of the structure, with CMP-Neu5Ac as sticks. The highlighted areas are sialylmotifs L, S, III, and VS (in orange, blue, red and green, respectively), as well as the key disulfide bridges as sticks. (b) Surface-view of the binding pocket (colors identical to (a)) and omit electron density map (2mFo-DFc) for CMP-Neu5Ac. (c) Close-up view of the CMP-Neu5Ac binding site (colors identical to (a)) and highlighted as sticks are residues involved in the binding. Please see main text for discussion of the alternative conformations A and B of residue Tyr354 (d) Interaction plot of the CMP-Neu5Ac; green: hydrogen bond; yellow: hydrophobic interaction; pink and purple: π-alkyl and amide π-stacking.

**Figure 2.**
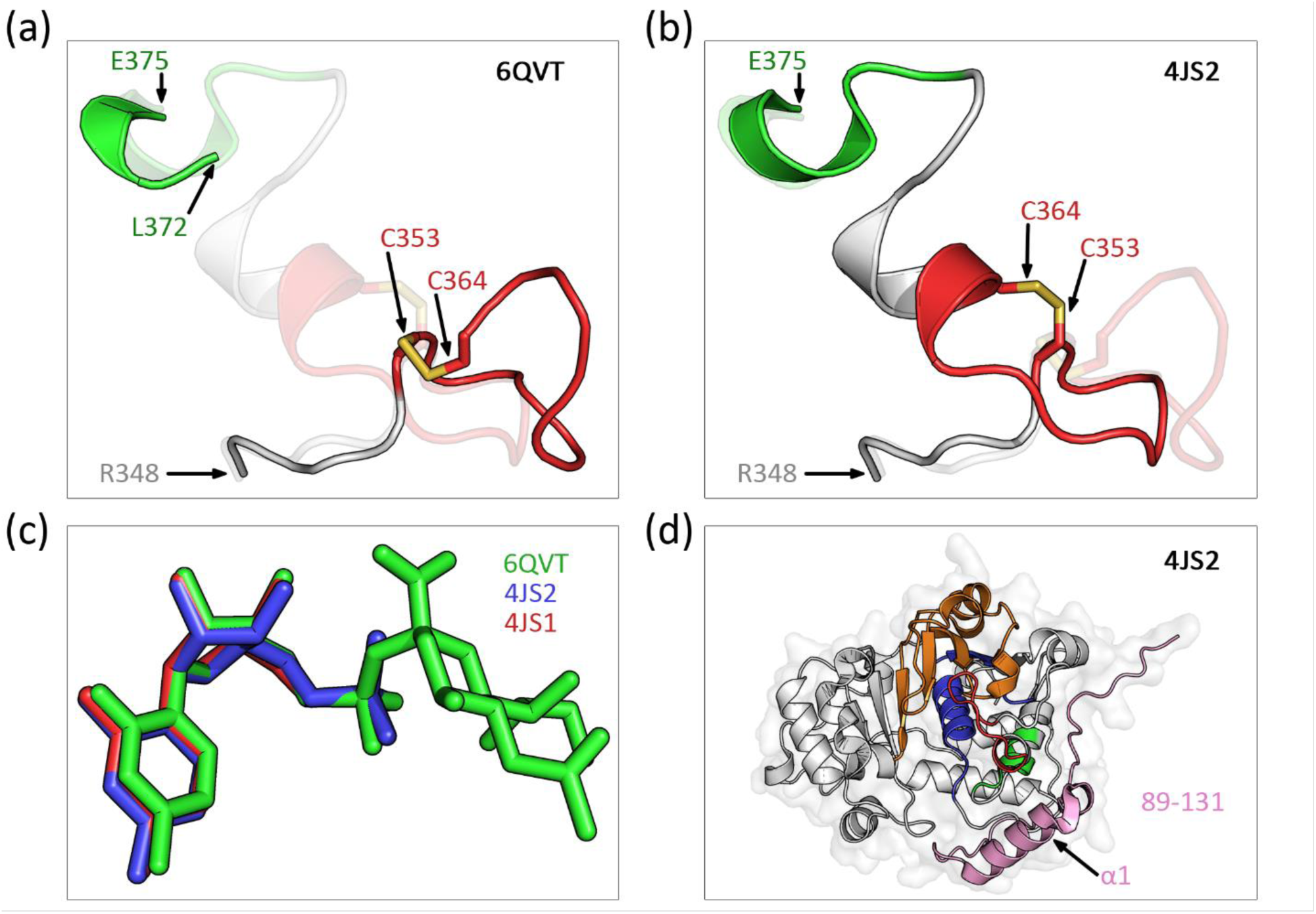
Differences between our structure and the previously published structures. (a, b) Close-up within the 348-375 region, with the disulfide bridge between C353 and C364 highlighted as sticks. For easier comparison, (a) shows our structure 6QVT with 4JS2 as a faded image, while (b) shows 4JS2 with 6QVT as a faded image. Colors are according to Fig. 1. (c) Superimposition of the three structures 6QVT (green, CMP-Neu5Ac), 4JS2 (blue, CMP) and 4JS1 (red, Cytidine) shows the similarity between positions of the ligands. (d) Difference in the overall construct range between 4JS1/4JS2 and our structure 6QVT: the region in pink is absent from our construct.

### 3.4. Comparison of the unliganded and CMP-Neu5Ac bound ST6Gal I structures

The two structures presented here, both overall and for the binding pocket, are very similar, with the exception of the movement of Tyr354. In the unliganded form, only conformation A of Tyr354 (see Figure 1c) is present. However, in the CMP-Neu5Ac bound structure, an alternate conformation B (refined at 48% occupancy) appears, showing the movement of this residue. It is interesting to note that Tyr354 in conformation A interacts with the CMP-Neu5Ac both at the phosphate and the Neu5Ac moiety. This suggests that the unliganded form with its conformation A is already adapted to interact with the substrate. In the CMP-Neu5Ac bound structure we see that Tyr354 conformation B is further away from the CMP-Neu5Ac. This in turn could be interpreted to represent a conformation of the protein preparing for the hydrolysis step. It is known that mutating Tyr354 results in a variant enzyme with low residual activity (Laroy *et al*., 2001). On the other hand, some other vertebrate ST6Gal I as well as ST6Gal II enzymes do have a histidine replacing the Tyr354. From our structure it can be envisaged that a histidine would maintain the hydrogen bond with the sugar portion of the donor substrate, but the distance towards the phosphate would extend to at least 4 Å. Yet other contacts would presumably hold the substrate amenable for catalysis.

### 3.5. Comparison of the present and previously published ST6Gal I structures

The protein for the previously published structures of ST6Gal I (Kuhn *et al*., 2013) was produced in human HEK293 cells, while in the present work protein was produced in *E. coli*. This explains the absence of glycosylation on Asn149 seen in the former structure. There is also difference in the amino acid residue range of the constructs used (see Table 1) as our structures are lacking an α- helix in position 100-121. The shorter construct was chosen based on comparison of initial expression and purification yields, and readily produced well-diffracting crystals. Pragmatically taken, our structure demonstrates that the omitted α-helix does not hamper ST6Gal I expression and folding. Furthermore, it also validates the biological relevance of the ligand-bound structure, since the cytidine monophosphate (CMP) or cytidine-phosphate moieties (incomplete substrates) bind identically in the structures by Kuhn *et al*. (2013) as the corresponding parts of the full-length substrate in our structure, i.e. the omitted α-helix is not critical for donor substrate binding.

The main differences observed between our liganded structure 6QVT and previous ST6Gal I structures are threefold. Firstly, in our structure the region 366-372, corresponding to family motif “d” and sialylmotif VS, is disordered. Possible explanations for this are that (i) binding of acceptor glycan is required for stabilizing this region, or that (ii) although the α-helix 100-121 does not affect the folding of the protein with regard to expression and structure determination, it may be required for stabilizing this region irrespective of acceptor glycan binding. Secondly, the disulfide bond C353-C364 is in a very different orientation, again allowing to hypothesize that the 353-370 region can move upon binding of the acceptor glycan. Thirdly, the donor substrate is bound with a multitude of hydrogen bonds and other interactions (Figure 1c&d) and the mobile loop is also contributing to substrate specificity, similarly as in ST3Gal I (Rao *et al*., 2009; Petit *et al*., 2015). Finally, CMP-Neu5Ac is bound with the side chain at C-5 of the sugar residue directed towards empty space at the surface of the protein. It is known that CMP-Neu5Ac sialic acid modified at the N-acyl position with a bulkier alkynyl group (Noel *et al*., 2018) can serve as the donor substrate both for α2,6-sialylation of N-glycans and α2,3-sialylation of O-glycans (Noel *et al*., 2017), and Sun *et al*. (2016) showed that even biotin at the N-acyl position can be tolerated and used to introduce improved labelling and identification of subsets of cell surface glycoconjugates. Our structure, with the CMP-Neu5Ac bound, explains the functionality of such modified unnatural sialic acid analogues with unique chemistry for metabolic labeling and other applications (Yarema, 2001), and may also encourage further attempts to produce inhibitors mimicking the natural donor substrate (Montgomery *et al*., 2017).

## ACKNOWLEDGEMENTS

We acknowledge the European Synchrotron Radiation Facility for provision of synchrotron radiation facilities and we would like to thank the local contacts for providing assistance in using the beamlines ID30A-3 and ID23-1 (proposals MX1850 and MX1933). This work was carried out with the support of Biocenter Oulu, <u>Structural Biology Core Facility,</u> University of Oulu, Finland. We thank Rik Wierenga for fruitful discussions and critical reading of the manuscript. This work has been funded by the Academy of Finland and University of Oulu, and also facilitated by University of Oulu short-term international research visits fund and the support provided jointly by the Institut Francais de Finlande, the Embassy of France in Finland, the French Ministry of Education, Higher Education and Research, and the Finnish Society of Science and Letters.

